# Endothelial TRPV4/Cx43 Signaling Complex Regulates Vasomotor Tone in Resistance Arteries

**DOI:** 10.1101/2024.07.25.604930

**Authors:** Pía C. Burboa, Pablo S. Gaete, Ping Shu, Priscila A. Araujo, Annie V. Beuve, Walter N. Durán, Jorge E. Contreras, Mauricio A. Lillo

**Affiliations:** Department of Pharmacology, Physiology and Neuroscience, Rutgers-New Jersey Medical School, Newark, NJ 07103, U.S.A; Department of Physiology and Membrane Biology, School of Medicine, University of California Davis, Davis, CA, U.S.A

**Keywords:** ECs, Caveolae, hyperpolarization, resting membrane potential

## Abstract

S-nitrosylation of Cx43 gap junction channels critically regulates communication between smooth muscle cells and endothelial cells. This posttranslational modification also induces the opening of undocked Cx43 hemichannels. However, its specific impact on vasomotor regulation remains unclear. Considering the role of endothelial TRPV4 channel activation in promoting vasodilation through nitric oxide (NO) production, we investigated the direct modulation of endothelial Cx43 hemichannels by TRPV4 channel activation. Using the proximity ligation assay, we identify that Cx43 and TRPV4 are found in close proximity in the endothelium of resistance arteries. In primary endothelial cell cultures from resistance arteries (ECs), GSK-induced TRPV4 activation enhances eNOS activity, increases NO production, and opens Cx43 hemichannels via direct S-nitrosylation. Notably, the elevated intracellular Ca^2+^ levels caused by TRPV4 activation were reduced by blocking Cx43 hemichannels. In ex vivo mesenteric arteries, inhibiting Cx43 hemichannels reduced endothelial hyperpolarization without affecting NO production in ECs, underscoring a critical role of TRPV4/Cx43 signaling in endothelial electrical behavior. We perturbed the proximity of Cx43/TRPV4 by disrupting lipid rafts in ECs using β-cyclodextrin. Under these conditions, hemichannel activity, Ca^2+^ influx, and endothelial hyperpolarization were blunted upon GSK stimulation. Intravital microscopy of mesenteric arterioles *in vivo* further demonstrated that inhibiting Cx43 hemichannels activity, NO production and disrupting endothelial integrity reduce TRPV4-induced relaxation. These findings underscore a new pivotal role of Cx43 hemichannel associated with TRPV4 signaling pathway in modulating endothelial electrical behavior and vasomotor tone regulation.

## INTRODUCTION

The vascular system is integral to maintaining cellular homeostasis by ensuring the delivery of oxygen and nutrients to each cell within the organism. At the forefront of vascular function is the endothelium, a specialized lining that interfaces with the blood vessel lumen. Endothelial cells (ECs) orchestrate critical signaling processes that regulate blood flow resistance, making them pivotal in preventing cardiovascular-related diseases.

Central to the control of blood flow resistance are the arterioles within the microcirculation^1–3^. These vessels, by modulating peripheral vascular resistance, play a crucial role in regulating arterial blood pressure and blood flow distribution. The intricate control of vessel diameter is mediated mainly through interactions among ECs, smooth muscle cells, and perivascular nerves^1–4^.

ECs exert tonic control over vasomotor tone primarily through the Ca^2+^-dependent production of vasodilator signals, nitric oxide (NO) and endothelium-dependent hyperpolarization (EDH) ^5–7^. While the precise biochemical mechanisms of EDH remain under investigation, evidence suggests its reliance on the activation of endothelial Ca^2+^-activated K^+^ channels, specifically small (SK_Ca_) and intermediate (IK_Ca_) conductance channels^5–7^. This hyperpolarization is transmitted to smooth muscle cells through connexin-formed gap junctions located at myoendothelial junctions ^5–7^.

Gap junction channels are formed by the docking of two hemichannels, each composed of hexameric single-membrane connexin subunits contributed by an adjacent cells. Gap junction channels allow direct cytoplasmic communication^8–10^. However, there is evidence that undocked hemichannels can also function independently at the plasma membrane, serving as a tightly regulated pathway for paracrine signaling or Ca^2+^ entry^11^. In myoendothelial junctions, S-nitrosylated Cx43 gap junction channels are crucial for regulating vascular diameter by facilitating the transmission of vital vasoactive molecules, such as Ca^2+^, IP_3_, NO, between endothelial and smooth muscle cells^12,13^. S-nitrosylation of Cx43 has been also shown to activate undocked hemichannels at the plasma membrane, influencing the membrane excitability of cardiomyocytes ^14–16^. Specifically, they play pivotal roles in cardiac action potentials and the influx of Ca^2+^ into cardiomyocytes under both normal conditions and during cardiac dysfunction^14,16,17^. Interestingly, the role of S-nitrosylated Cx43 hemichannels in normal vascular physiological processes like as in the EDH generation in ECs, as well as their role in regulating vasomotor tone, remains largely unexplored.

TRPV4 activation, whether induced by endothelium-dependent vasodilators or shear stress, plays a crucial role in orchestrating endothelial Ca^2+^ signaling, leading to the generation of NO and mainly activation of the EDH ^6,18–22^. Sonkusare et al ^6^. demonstrated the significance of TRPV4 in endothelial Ca^2+^ influx, particularly in promoting activation of Ca^2+^-activated potassium channels (K_Ca_), which are essential for EDH-mediated vascular dilation. Consistent with this, studies on TRPV4 knockout mice have shown impaired acetylcholine (ACh)-induced NO/hyperpolarization and arteriolar dilation, highlighting the indispensable role of TRPV4 in maintaining vascular homeostasis ^6,18–22^. Interestingly, Cx43 and TRPV4 channels are compartmentalized within the myoendothelial projections to facilitate effective communication between ECs and smooth muscle in arterioles^23^. Therefore, we aim to investigate the potential relationship between TRPV4-induced vasodilation via NO production, S-nitrosylation and opening of Cx43 hemichannels, and their role in vasomotor function.

We discovered that TRPV4 promotes the S-nitrosylation of Cx43 hemichannels in ECs. Furthermore, there is a close spatial proximity between TRPV4 and Cx43 proteins, predominantly observed in the endothelium of resistance arteries. The TRPV4/Cx43 complex plays a pivotal role in regulating vasomotor tone by mainly influencing endothelial electrical activity.

## RESULTS

### TRPV4 associates closely with Cx43, and its Activation promotes S-nitrosylation and opening of Cx43 Hemichannels

Figure 1 displays different mouse mesenteric arterioles where the main interaction between Cx43 and TRPV4 is observed in the endothelial layer compared to the smooth muscle cells (red stain). A few dots are visible in the smooth muscle cells (SMC), while strong interactions are observed near the endothelium (EC) and smooth muscle cells within the lamina (green), where the myoendothelial junction is situated. However, as depicted in several insets of the displayed vessels, there is also a significant amount of signal around endothelium close to the lumen (*). This indicates that the Cx43/TRPV4 interaction is not exclusively confined to the myoendothelial junctions and is constitutively present in the ECs compared to the smooth muscle cells.

**Figure 1.**
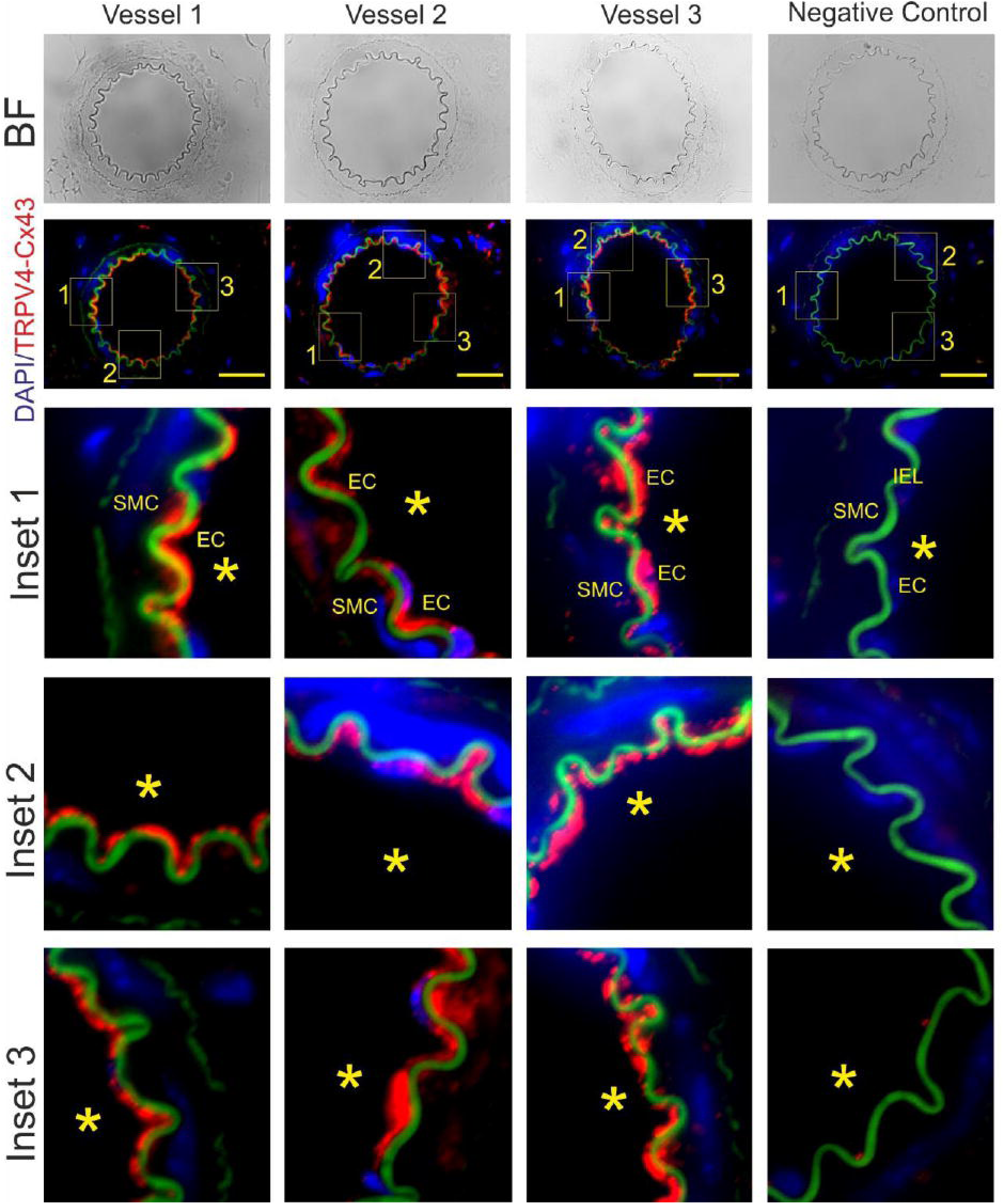
TRPV4/Cx43 complex is primarily observed within the endothelial cell layer of resistance arteries. Proximity Ligation Assay (PLA) was utilized to illustrate the spatial co-localization between TRPV4 and Cx43. Notably, TRPV4 and Cx43 exhibit predominant distribution within the endothelial layer (EC) compared to the smooth muscle cells (SMC). Controls were conducted to validate the specificity of PLA analysis; wherein primary antibodies were omitted as a negative control. Asterisks indicate the luminal region of the vessels, while the green, fluorescent signal corresponds to the internal elastic lamina (IEL). (BF: Brightfield, Scale bar: 80 µm)

Shear stress activates endothelial TRPV4, leading to the production of NO^5–7^. We previously showed that NO enhances the activity of Cx43 hemichannels via S-nitrosylation in a heterologous expression system^14^. Building upon these findings, we sought to determine whether a specific and potent TRPV4 activator, GSK 1016790A (10 nM), induces the S-nitrosylation of Cx43 and the opening of hemichannels.

We first observed that GSK-induced TRPV4 activation enhances the phosphorylation of eNOS at S1177 while simultaneously decreasing the phosphorylation of eNOS at T495 (Figure 2A) in primary cultures of ECs from mesenteric arterioles. This posttranslational modifications in eNOS phosphorylation confirms the enhancement of eNOS activity via TRPV4 activity. Overall, we observed that GSK increases the S-nitrosylation of protein levels in ECs (Figure 2A).

**Figure 2.**
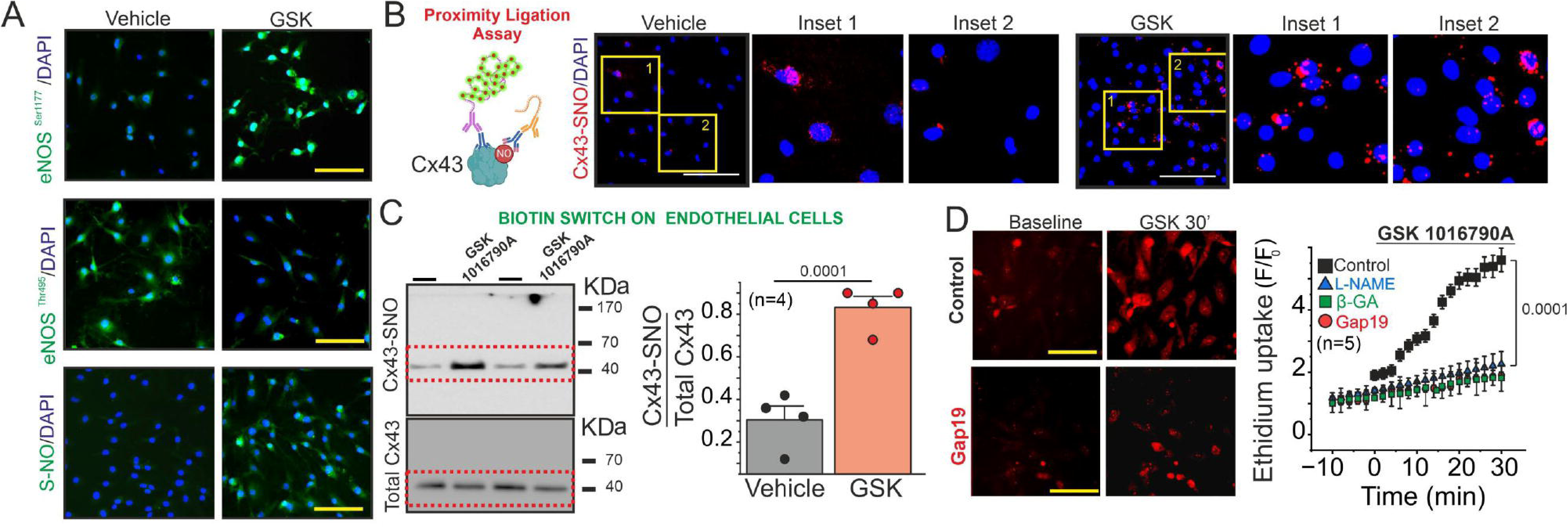
TRPV4 activation promote S-nitrosylated Cx43 hemichannels activity by NO production. **A**) Representative immunofluorescence microscopy images illustrating the detection of phosphorylation status at Serine 177 and Threonine 495 residues of endothelial nitric oxide synthase (eNOS), alongside the assessment of protein levels of S-nitrosylated proteins, in vehicle conditions and upon stimulation with 10 nM GSK 1016790A on primary cultures of ECs. **B**) Representative images from proximity ligation assay demonstrating Cx43 S-nitrosylation levels in both control and 10 nM GSK 1016790A-stimulated conditions. **C)** Western blot analysis (left) and quantification (right) of S-Nitrosylated Cx43. The S-nitrosylated Cx43 levels were expressed as fold change relative to total Cx43 protein levels per sample. Comparisons between groups were made using Student’s t test. **D)** Time course analysis to measure the uptake of 5 µM ethidium, assessing hemichannel activity upon 10 nM GSK 1016790A and under pre-treatment of 100 µM L-NAME, 50 µM glycyrrhetinic acid (β-GA), and 50 µM Gap19. Statistical comparisons between the time course curves were performed using one-way ANOVA followed by Tukey post hoc tests. The total number of mice used in the experiments is indicated in parentheses. (scale bar: 80 um)

Next, we tested whether TRPV4 activation induces Cx43 S-nitrosylation using a Proximity Ligation Assay (PLA) and Biotin Switch approach, as we previously performed in cardiac tissue^14,16^. We found that under control conditions, there are few Cx43 S-nitrosylated proteins in in ECs detected by PLA experiments. However, administration of 10 nM GSK 1016790A, caused a substantial increase in S-nitrosylated Cx43 signal compared to control conditions (Figure 2B). Similar results were obtained using the biotin switch assay approach. Under basal conditions, ECs displayed low levels of S-nitrosylated Cx43 protein, which was significantly elevated when the cells treated with the TRPV4 agonist, 10 nM GSK 1016790A. (Figure 2C, Cx43-SNO/ Total Cx43: Baseline = 0.305 ± 0.065; GSK = 0.8325 ± 0.05218). These findings demonstrate that activation of TRPV4 channels stimulate S-nitrosylation of Cx43 in ECs via eNOS activation.

To assess the functional impact of TRPV4-induced Cx43 S-nitrosylation, we investigated whether the TRPV4 agonist, enhances Cx43 hemichannel activity in ECs by measuring ethidium (Et) uptake at 5 µM. TRPV4-activation induced a strongly ethidium uptake in ECs during the time (Figure 2D). Either inhibition of NO production with 100 µM L-NAME or the blockade of Cx43 hemichannels with 50 µM β-glycyrrhetinic acid (β-GA) or 50 µM Gap19-TAT, a specific Cx43 hemichannel blocker ^15,16,24,25^, significantly reduced dye uptake increases. (Figure 2D). These findings suggest that activation of TRPV4 channels promotes the opening of Cx43 hemichannels in ECs from resistance arteries through a NO production and Cx43 S-nitrosylation.

To further support that activation of TRPV4 channels enhances Cx43 hemichannel activity via direct S-nitrosylation, we assessed ethidium uptake in a heterologous expression system (HeLa cells) co-transfected with TRPV4 and either Cx43 or a Cx43 mutant lacking the S-nitrosylation site (Cx43 C271S) ^14^. HeLa cells expressing TRPV4 or Cx43 alone showed a slight increase in Et uptake rates compared with control EGFP-transfected cells (**Supplemental Figure 2**). However, co-expression of TRPV4 with Cx43 led to a robust increase in ethidium uptake, which was slightly increased in the presence of GSK 1016790A (**Supplemental Figure 2**). Consistently, HeLa cells expressing Cx43 C271S displayed a reduced Et uptake rate, similar to that observed in EGFP-expressing cells, without affecting Cx43 expression or distribution (**Supplemental Figure 3**). More importantly, the C271S mutation prevented the potentiation of Et uptake induced by the co-expression of TRPV4 and application of GSK 1016790A (**Supplemental Figure 2**). These results indicate that TRPV4 expression enhances Cx43 hemichannel activity via Cx43 S-nitrosylation at the C271 residue.

### Endothelial Ca^2+^ influx induced by TRPV4 channel activation is mediated by Cx43 hemichannels

TRPV4 activation plays a crucial role as a mediator in endothelial Ca^2+^ influx. Our research, along with others, demonstrated that Cx43 hemichannels contribute to Ca^2+^ increases in both physiological and pathophysiological conditions in ventricular cardiomyocytes^14–16^. However, it remains unclear whether the signaling interaction between TRPV4 and Cx43 hemichannels mediates Ca^2+^ influx in ECs.

We assessed global Ca^2+^ increases in ECs using Fluo-4 Ca^2+^ indicator ^4,7,26^. We observed a biphasic response to 10 nM GSK 1016790A, characterized by an initial peak within 10 seconds followed by a sustained plateau after 10 seconds. Both phases of the Ca^2+^ response were diminished upon inhibition of Cx43 hemichannels using 50 µM Gap19-TAT (Figure 3A-B, Peak Gap19-TAT: 1.07 ± 0.28, A.U; Peak Gap19-TAT Plateau: 0.038 ± 0.0181, A.U).

**Figure 3.**
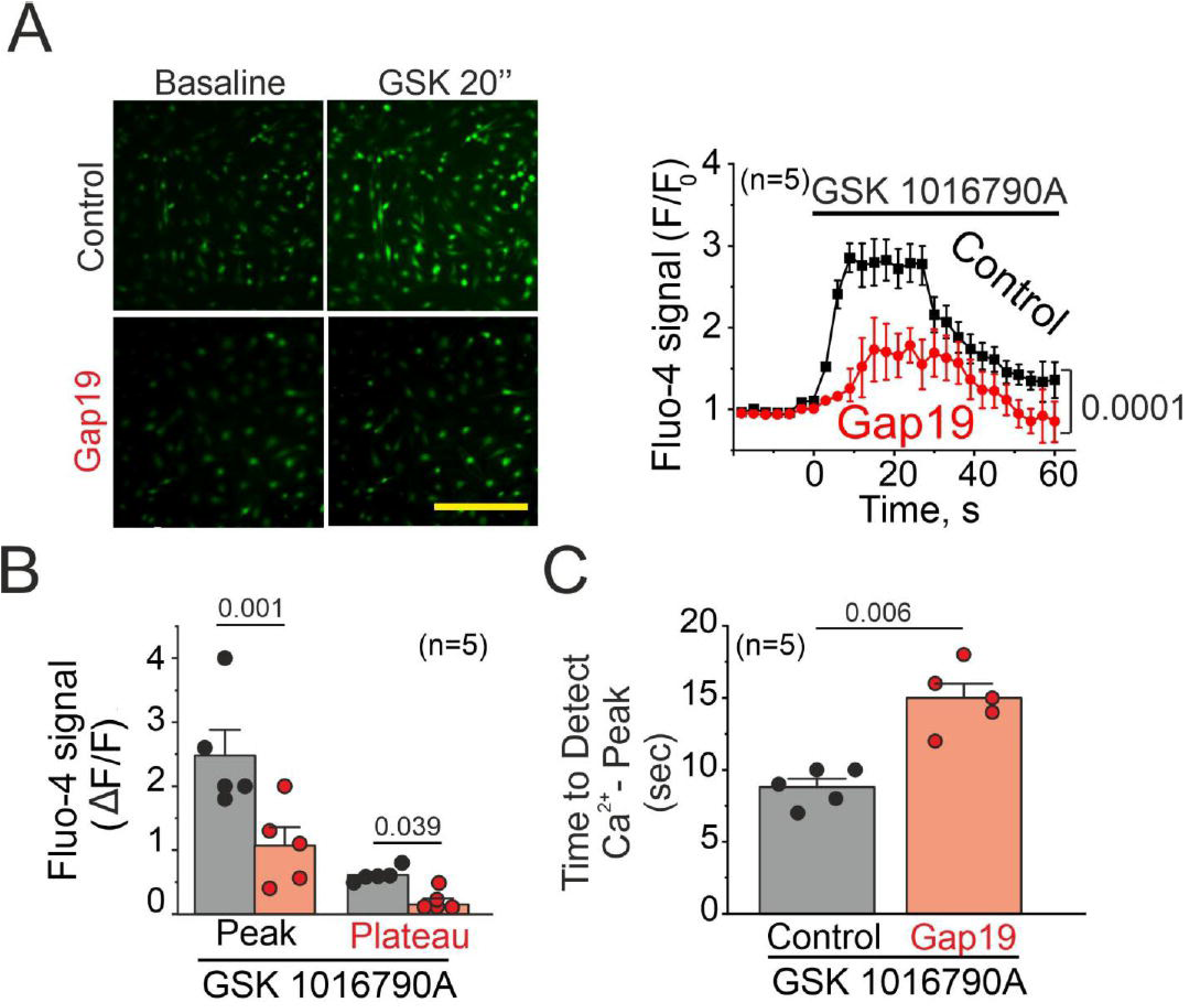
TRPV4/Cx43 hemichannels mediate Ca^2+^ increases in endothelial cells in response to TRPV4 activation. **A**) Left: Representative imaging captures the endothelial cell responses before and after exposure to 10 nM GSK 1016790A stimulation, both under control conditions and in the presence of 50 uM Gap19. Right: Time course analysis illustrates the fluctuations in Fluo-4 signaling subsequent to GSK stimulation. **B)** Peak Ca^2+^ increases observed in A. Comparisons between groups were made using Student’s t test. **C)** Analysis conducted to assess the time required to detect the Ca^2+^ peak under control conditions and in the presence of 50 µM Gap19 upon GSK simulation. Comparisons between groups were made using Student’s t test. The total number of mice used in the experiments is indicated in parentheses. (scale bar: 80 um)

Furthermore, activation of TRPV4 elicited a faster increase in Ca^2+^ levels compared to cells with inhibited Cx43 hemichannels activity (Figure 3C, Time to detect Ca^2+^ peak, Control: 8.8 ± 0.58 min, Gap19-TAT: 15 ± 1.1 sec). These findings provide compelling evidence that endothelial Ca^2+^ increases induced by TRPV4 activation are mediated by endothelial Cx43 hemichannels.

### Cx43 hemichannels facilitate endothelial hyperpolarization following TRPV4 activation

Our findings depicted in Figure 3 strongly suggest a novel role of Cx43 hemichannels in regulating Ca^2+^ influx upon TRPV4 activation. Given that TRPV4 plays a crucial role in NO and EDH generation^6,18–22^, we sought to investigate whether Cx43 hemichannels are also pivotal in NO and EDH generation. ECs were exposed to 10 µM DAF-2DA, a fluorescent probe used for detecting NO ^13^. Control conditions exhibited a rapid increase in NO production following stimulation with 10 nM GSK 1016790A. Similarly, cells pre-treated with 50 µM Gap19-TAT showed a comparable rate of NO generation **(Supplemental Figure 4)**. To confirm that NO synthesis is indeed induced by TRPV4 activation, ECs were pre-treated with 100 µM L-NAME for 90 minutes to block NO production. Under these conditions, there was a significant reduction in NO generation signaling **(Supplemental Figure 4)**. These data suggest that inhibition of Cx43 hemichannels does not affect endothelial NO synthesis upon TRPV4 activation.

Next, we investigated whether TRPV4/Cx43 hemichannel signaling is involved in endothelial hyperpolarization. To address this gap in knowledge, we utilized intact *en face* mouse mesenteric arteries to directly assess the involvement of Cx43 hemichannels in endothelial hyperpolarization upon TRPV4 activation.

Mesenteric arteries ranging from 3rd to 4th order, with inner diameters between 150-200 μm, were longitudinally sectioned to expose the endothelial layer. Cell labeling using FITC-70KDa-Dextran in the recording pipette facilitated visualization of the cells (Figure 4A). Blocking Cx43 hemichannels resulted in a significant reduction in endothelial hyperpolarization peak induced by 10 nM GSK 1016790A (Figure 4A, Control: –15.26 ± 0.94 mV, Gap19-TAT: – 5.88 mV ± 0.83). To determine that the endothelial hyperpolarization induced by GSK is mediated via K_Ca_ potassium channels, we employed 200 nM charybdotoxin, a specific blocker for intermediate K_Ca_ channels, and 300 nM of apamin, a specific blocker for small K_Ca_ channels^27^. Under these experimental conditions, we observed a significant reduction in endothelial hyperpolarization peak (Figure 4A **Value:** –1.62 ± 0.29 mV). These data demonstrate that Cx43 hemichannels are involved in endothelial hyperpolarization upon TRPV4 activation.

**Figure 4.**
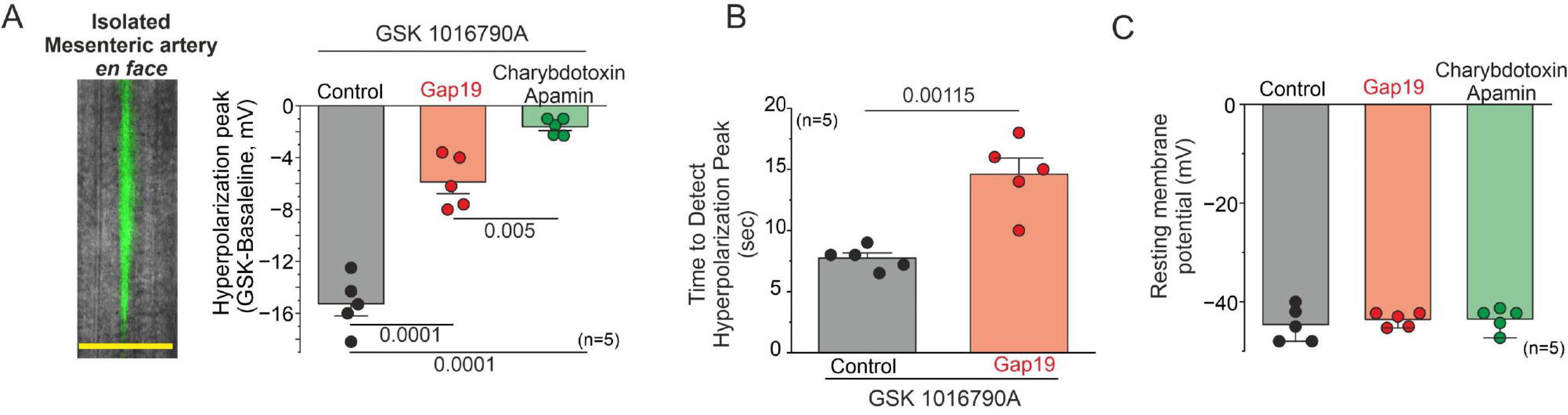
Cx43 hemichannels mediate endothelium hyperpolarization upon TRPV4 activation in intact endothelium. **A**) Left: Representative *en face* imaging of an arteriole showing ECs detected using FITC-Dextran 3KDa in the pipette solution to measure resting membrane potential changes. Right: Hyperpolarization peaks were induced by TRPV4 activation in intact ECs from mesenteric arteries under control conditions, 50 µM Gap19, and KCa channel blockers (300 nM charybdotoxin and 200 nM apamin). Group comparisons were made using nested ANOVA. **B)** Time to detect hyperpolarization peaks in ECs treated under control and 50 µM Gap19 conditions following stimulation with 10 nM GSK 1016790A. Group comparisons were made using Student’s t-test. **C)** Resting membrane potential of arteries under control conditions, with 50 µM Gap19, and with 300 nM charybdotoxin and 200 nM apamin. Group comparisons were made using nested ANOVA. The total number of mice used in the experiments is indicated in parentheses. (scale bar: 20 um)

Additionally, differences were observed between control conditions and cells treated with Gap19-TAT in the time required to detect the maximum hyperpolarization peak induced by GSK (Figure 4B, Control: 7.74 ± 0.42 seconds, Gap19-TAT: 14.6 ± 1.32 seconds), indicating that blocking Cx43 hemichannels promotes a delayed response in both endothelial Ca^2+^ increases (Figure 3) and endothelial hyperpolarization (Figure 4) upon TRPV4 activation. Prior to stimulation with GSK, ECs under control conditions, pre-treated with Gap19 and charybdotoxin/apamin, exhibited a similar resting membrane potential of approximately –45 to – 40 mV. (Figure 4C).

### Disruption of eNOS/TRPV4/Cx43 proximity decreases Ca^2+^ influx, and endothelial hyperpolarization

We conducted a proximity ligation assay in our primary culture of ECs to examine the proximity between Cx43, TRPV4 and eNOS. We observed a robust proximity between these proteins (Figure 5A). Next, we disrupted the compartmentalization of these proteins by employing 5 mM β-cyclodextrin to deplete the cholesterol-rich endothelial lipid rafts^26,28^. Under these conditions, the proximity between Cx43/eNOS/TRPV4 was significantly diminished.

**Figure 5.**
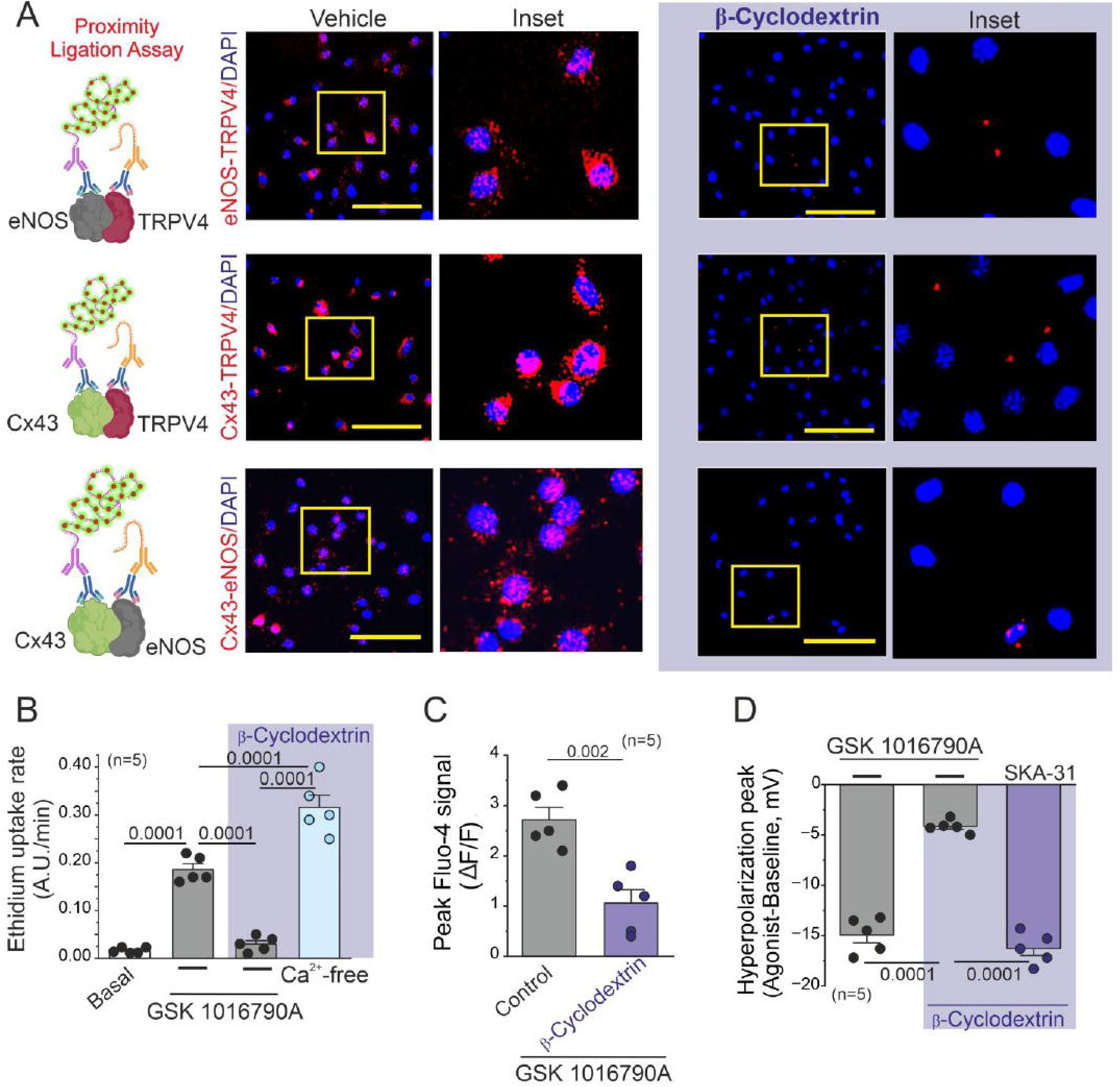
Critical Role of eNOS/Cx43/TRPV4 proximity in regulating endothelial Ca^2+^ increase and electrical behavior. **A**) Representative imaging of proximity ligation assay (PLA) in control conditions and in ECs treated with 5 mM β-Cyclodextrin to assess the spatial proximity between eNOS-TRPV4, Cx43-TRPV4, and Cx43-eNOS. Strong interactions between these proteins are indicated by red dots, while complete disruption of the PLA signal is observed with 5 mM β-Cyclodextrin treatment. B) Ethidium bromide uptake rate of ECs following stimulation with 10 nM GSK 1016790A under control conditions, extracellular zero Ca^2+^, and β-Cyclodextrin treatment. Group comparisons were conducted using nested ANOVA. C) Changes in intracellular calcium peak increases under control and cyclodextrin conditions in cells treated with 10 nM GSK 1016790A. D) Hyperpolarization peak induced by 10 nM GSK 1016790A under control and cyclodextrin conditions. To assess the activity of KCa channels under β-Cyclodextrin conditions, we used 10 µM SKA-3. Group comparisons were made using nested ANOVA. The total number of mice used in the experiments is indicated in parentheses. (scale bar: 80 um)

Under conditions of 5 mM β-cyclodextrin, we performed a 5 µM Ethidium dye uptake assay to assess Cx hemichannel activity (Figure 5B). We observed a pronounced reduction in dye uptake in ECs upon 10 nM GSK1016790A stimulation under β-cyclodextrin conditions. (Basal: 0.002 ± 6.78E-4, A.U/min, GSK: 0.186 ± 0.012, A.U/min; GSK + β-cyclodextrin: 0.03 ± 0.007, A.U/min). To ascertain whether Cx hemichannel activity is impacted by β-cyclodextrin treatment, we performed tests under conditions of zero extracellular Ca^2+^, which is an effective strategy for assessing hemichannel dye uptake^13,29,30^. We observed an increase in dye uptake rate (β-Cyclodextrin + Ca^2+^-Free: 0.316 ± 0.025, A.U/min), suggesting that β-cyclodextrin treatment does not influence Cx hemichannel function in ECs.

Following our findings that Cx43 hemichannels are involved in Ca^2+^ influx (Figure 3), we investigated whether disruption of TRPV4/Cx43 proximity also affected Ca^2+^ increases upon TRPV4 activation. Under the same aforementioned conditions, ECs loaded with 10 µM Fluo-4 displayed a significant reduction in Ca^2+^ increases (Figure 5C, Control: 2.72 ± 0.24 A.U, β-Cyclodextrin: 1.1 ± 0.19 A.U.), as well as diminished endothelial hyperpolarization upon 10 nM GSK 1016790A stimulation (Figure 5D, Control: –14.94 ± 0.78 mV, β-Cyclodextrin: –4.16 ± 0.28 mV). As other research groups have demonstrated the co-localization of KCa channels and TRPV4 within endothelial microdomains ^6,31–33^, we investigated whether β-cyclodextrin affects the activity of K_Ca_ potassium currents. Under cyclodextrin conditions, we stimulated ECs with 10 µM SKA-31, an activator of IK_Ca_ and SK_Ca_ channels known to promote endothelial hyperpolarization^34–36^. Remarkably, under these conditions, the peak endothelial hyperpolarization remained comparable to control values (Figure 4A), suggesting that cyclodextrin treatment did not influence K_Ca_ channel activity (SKA-31 + Cyclodextrin: –16.28 ± 0.7 mV).

Overall, these results underscore the pivotal role of the proximity and signaling interactions between eNOS, TRPV4, and Cx43 hemichannels in modulating endothelial electrical vascular function.

### Cx43 hemichannels contribute to vascular response in mesenteric arterioles *in vivo* following TRPV4 activation

We subsequently investigated the potential role of TRPV4/Cx43 signaling activation in the regulation of vasomotor tone in intact mesenteric vessels *in vivo*, utilizing intravital microscopy To enhance visualization of arterioles within the mesenteric vascular bed, we administered FITC-70KDa-Dextran via retroorbital injection (Figure 6A-B).

**Figure 6.**
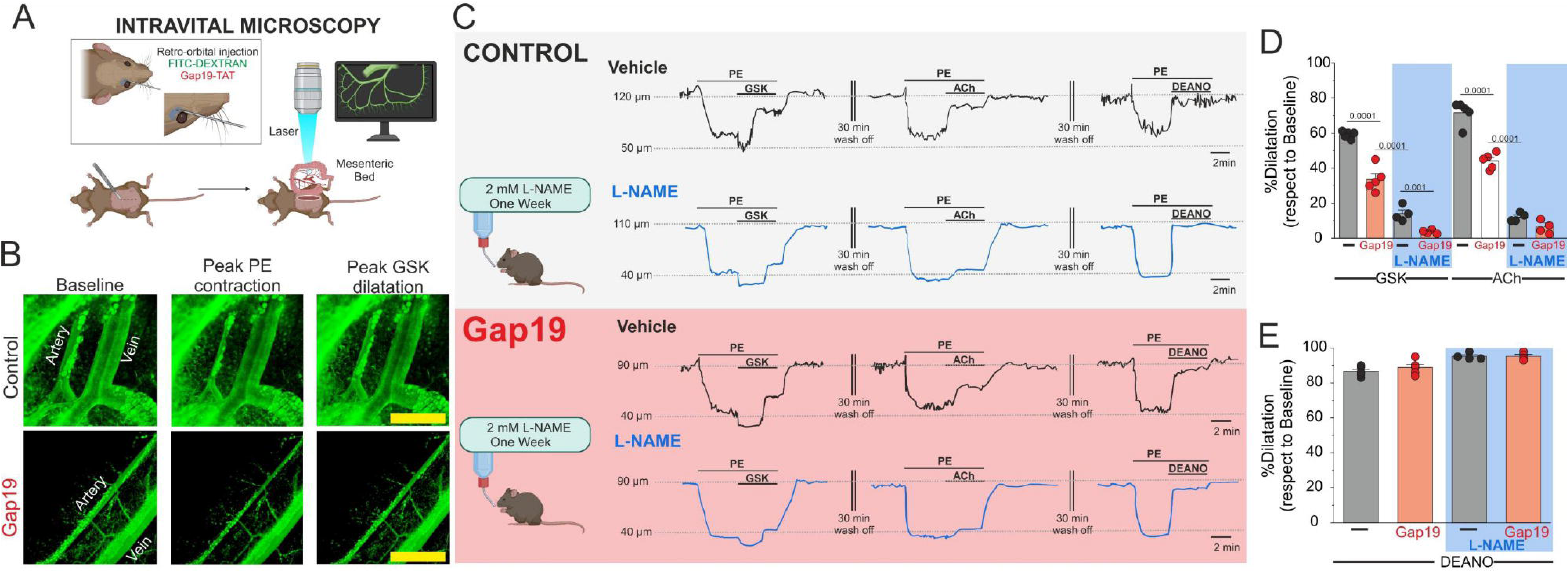
Endothelial Cx43 hemichannels mediate vasomotor responses in mesenteric arteries in vivo. **A**) Schematic illustrating intravital microscopy approaches. To enhance vessel visualization, 50 µg/kg FITC-dextran (70 kDa) and 10 µg/kg Gap19, to block Cx43 hemichannel activity, were administered via retroorbital injection. **B)** Representative imaging of the mesenteric bed at baseline, during the phenylephrine (PE) peak, and at the dilation peak in control and 50 µM Gap19 conditions. **C)** Representative traces of vasomotor responses induced by 100 nM GSK 1016790A, 10 µM PE, 10 µM ACh, and 10 µM NO donor DEANO in control, 50 µM Gap19 and L-NAME conditions. **D)** Percentage of relaxation in vessels pre-contracted with 10 µM PE upon GSK, 10 µM ACh. Group comparisons were made using Student’s t-test. The total number of mice used in the experiments is indicated in parentheses. **E)** Percentage of relaxation in vessels-precontracted with 10 µM PE upon 10 µM DEANO in control and Gap19-treated conditions. Group comparisons were made using Student’s t-test. The total number of mice used in the experiments is indicated in parentheses. (scale bar: 500 um).

We tested the mesenteric vascular relaxation in arteries pre-contracted with 10 µM Phenylephrine (PE). Under these conditions, 100 nM GSK induced a relaxation of 60-70% under control conditions while arterioles from mice pre-treated with 10 µg/kg of Gap19-TAT via retroorbital injection^14,16,37^ showed around 30 % relaxation (Figure 6C **and 6D**: % Relaxation: Control: 59.04 ± 0.91 %, Gap19-TAT: 33.6 ± 3.2 %). To avoid the biphasic vasomotion responses triggered by TRPV4 activation upon PE stimulation *in vivo*, and to investigate mainly endothelial vascular effect, we employed 10 µM acetylcholine (ACh), an endothelium-dependent vasodilator known to activate TRPV4 channels via PKC in ECs^38^. Under these conditions, we observed 71.6 ± 2.9 % relaxation compared to vessels treated with Gap19-TAT, where the % relaxation was 44.06 ± 1.9 (Figure 6C-D). Finally, under both control and Gap19-TAT conditions, 10 µM diethylamine NONOate (DEANO, nitric oxide donor) induced almost complete relaxation (Figure 6C-E, Control: 86.8 ± 1.28, Gap19-TAT: 88.8 ± 1.88). These results confirm that the smooth muscle NO-GCs-PKG signaling pathway remains unaffected in mice treated with Gap19-TAT.

Next, to test the role of Cx43 hemichannels on smooth muscle machinery response, we conducted a concentration-response curve of phenylephrine (PE), an α-adrenergic receptor that promotes smooth muscle cell contraction and depolarization ^39,40^. We did not observe differences in vascular contraction *in vivo* between vessels treated with and without Gap19-TAT (**Supplemental Figure 5A**). Similarly, to assess potential dysfunction in smooth muscle electrical properties due to Cx43 hemichannel blockade, we measured changes in the smooth muscle membrane potential in isolated mesenteric arteries following PE stimulation (**Supplemental Figure 5B**). Neither control nor arteries pre-treated with Gap19-TAT showed alterations in muscle depolarization induced by 10 µM PE. These results indicate that Cx43 hemichannels are not involved in smooth muscle functional activity.

Next, we investigated the impact of nitric oxide (NO) production *in vivo* within arterioles following GSK stimulation. To achieve this, we administered a 5 mM L-NAME solution in water to the mice for one week to suppress NO production to the greatest extent possible, as previously published ^14^. Under these control conditions, including Gap19-TAT treatment, the relaxation induced by GSK or ACh was significantly attenuated. (Figure 6D, PE + GSK + Gap19 + L-NAME: 3.5 ± 0.64; PE+ACh+Gap19+L-NAME: 9.25 ± 2.17), while the response to DEA-NO was unaffected (Fig. 6E).

To confirm the involvement of the endothelium, we compromised the endothelial layer by subjecting arterioles to prolonged laser/light exposure, using procedures conducted by others^41–44^. Arterioles treated with GSK upon PE stimulation displayed primarily a contraction component, similar to the results observed under L-NAME conditions **(Supplemental Figure 6A-B and Figure 6).** Additionally, ACh-induced relaxation was completely attenuated **(Supplemental Figure 6A-B).** To test whether the vascular response remained unaltered, we applied the NO donor, 10 µM DEANO. Under NO donor treatment, arterioles returned to initial baseline tone **(Supplemental Figure 6A-B).**

## DISCUSSION

Here we identified a functional interaction between Cx43 hemichannels and TRPV4 channel activation in regulating vascular resistance. We demonstrate that: **(1)** the TRPV4/Cx43 complex is predominantly in spatial proximity in the ECs of intact resistance vessels. **(2)** The signaling pathways involving endothelial TRPV4/Cx43 hemichannels represent a pivotal nexus in the endothelium Ca^2+^ increases. **(3)** NO plays a crucial role in smooth muscle relaxation not only through the NO-PKG signaling pathway but also via the endothelial nitrosylation of Cx43, thereby contributing to endothelial electrical hyperpolarization.

Although the compartmentalization of TRPV4 and Cx43 proteins has been previously described ^23^, our observations indicate that this compartmentalization predominantly occurs within ECs along the lamina, rather than being exclusively confined to myoendothelial junction projections. This suggests a potential interaction between TRPV4 channels and undocked Cx43 hemichannels. In other cell types, such as lens epithelium, TRPV4 activators have been shown to enhance ion channel conductance similar to Cx43 hemichannel^45^. Similarly, mechanical stimulation in astrocytes promotes a signaling pathway involving TRPV4 activation, which correlates Cx43 hemichannel opening and gliotransmitter release^25^. We consistently found a functional interaction between TRPV4 channel activation and opening of Cx43 hemichannels in ECs, which required production of NO and direct S-nitrosylation of Cx43. This functional interaction was further confirmed employing HeLa cells, as an a heterologous expression system, where the co-expression of Cx43 and TRPV4 increases hemichannel activity, as assessed by ethidium uptake. We previously demonstrated that Cx43 hemichannels activation required direct S-nitrosylation of cysteine at position C271^12,14^. As expected, Cx43 lacking cysteine 271, was insensitive to co-expression of TRPV4 channels, with not or little ethidium uptake detected. This indicates that TRPV4 channels-induced opening of Cx43 hemichannels required S-nitrosylation of Cx43.

Recent studies, including our own, have highlighted the potential physiological roles of Cx43 hemichannels. For instance, these studies have linked Cx43 hemichannels to endothelial cell migration^46^, cardiac action potentials and Ca^2+^ influx in cardiomyocytes during cardiac function ^14,17,47,48^. Additionally, Cx43 hemichannels have been implicated in D-serine release by astrocytes, and redox protection through the permeation of antioxidant metabolites, such as glutathione, in lens epithelium^49,50^. However, the role of Cx43 hemichannels in the regulation of vasomotor tone had not been previously examined. Our current work indicates that TRPV4 activation promotes NO production by enhancing eNOS activity, and consequently S-nitrosylation and opening of Cx43 hemichannels in ECs. When we blocked Cx43 hemichannels using Gap19-TAT, a mimetic peptide that does not block Cx43 gap junction channels^4,14,16,51–56^ or potassium conductance^14^, we observed a decrease on endothelium hyperpolarization, but no changes NO production. Furthermore, using Gap19, we observed a delay in endothelial Ca^2+^ increases and a similar delay in endothelial hyperpolarization in cells treated with TRPV4 activator. These findings suggest that TRPV4, NO and Cx43 hemichannels are part of the same signaling pathways and are critical mediators in Ca^2+^-induced EDH signaling.

Our results indicate that the TRPV4/Cx43 complex is essential for promoting endothelial hyperpolarization via TRPV4 activation. The TRPV4/Cx43 complex is in close proximity and disrupting this complex affected Ca^2+^ influx and endothelial hyperpolarization. Although some studies have indicated that TRPV4 activity in ECs depends on the presence of cholesterol^57^, or that cyclodextrin blocks the pores of connexin hemichannels in isoforms not expressed in ECs, such as Cx26 and Cx32^58^, our data derived from ECs suggest otherwise. Specifically, our findings indicate that hemichannel activity is still detectable under cyclodextrin conditions and zero extracellular Ca^2+^. Additionally, to exclude the potential effect of cyclodextrin on TRPV4/K_Ca_ signaling activity, we used a K_Ca_ activator, detecting similar endothelial hyperpolarization as with TRPV4 activation.

TRPV4 activation in microcirculation is linked to smooth muscle or endothelial hyperpolarization^18,23,32,59^, though reports indicate that TRPV4 activation in smooth muscle can also lead to contraction in arteries ^32^ and in other territories^60,61^. This variation in resistance arteries likely depends on the number of active TRPV4 channels, Ca^2+^ influx, and the proximity/interaction between critical proteins involved in smooth muscle contraction or dilatation^32,59^. For instance, recent findings revealed that stimulation of α1-adrenergic receptors (α1AR) activates TRPV4 channels in smooth muscle cells (TRPV4_SMC_) via protein kinase Cα (PKCα) signaling, significantly contributing to membrane depolarization and vasoconstriction and an increase in blood pressure ^32^. Conversely, TRPV4_SMC_ channel activity induced by intraluminal pressure counteracts vasoconstriction by activating Ca^2+^-sensitive K^+^ (BK) channels, demonstrating that there are functionally distinct pools of TRPV4_SMC_ channels^32^. Our experiments show that *in vivo* topical TRPV4 activator upon PE stimulation produces a biphasic response: an initial contraction followed by vasodilation. The initial contraction may be due to TRPV4_SMC-_PKCα signaling or the release of vasoconstrictor molecules (e.g. ATP) from smooth muscle cells, adipocytes or sympathetic or perivascular nerves around mesenteric arteries, which also modulate vascular activity ^62^. Interestingly, this *in vivo* vasodilation upon TRPV4 activation is reduced in the presence of Gap19, maintaining a high level of contraction similar to responses observed when the endothelium is compromised, consistent with the enhancement of α-adrenergic signaling via TRPV4 channels in smooth muscle ^32^. The precise role of TRPV4 channels in *in vivo* vascular models remains unclear, as their function may depend on their specific membrane nanodomain locations and associations with proteins that cause either depolarization or hyperpolarization of the membrane, leading to contraction or dilation, as has been postulated by other studies involving isolated cells, isolated vessels, or electrophysiological analyses of TRPV4^18,23,32,59^. Further experiments investigating the *in vivo* role of TRPV4, involving nerves, adipocytes, ECs, muscle cells, and other cell types in peripheral resistance, are essential to better understand the physiological role of TRPV4/Cx43 hemichannel signaling in controlling vascular responses.

Our data indicate that NO production is essential for activating endothelial Cx43 hemichannels, inducing endothelial hyperpolarization and vascular relaxation. The role of NO as a candidate for endothelium hyperpolarization factor remains controversial^63–66^. However, our findings align with previous studies showing that eNOS knockout reduces hyperpolarization in response to endothelium-dependent vasodilators, indicating that NO production is a critical component of endothelial electrical activity (i.e., EDH)^66^. Besides, it has been previously demonstrated that NO blockers require a more extended period to effectively inhibit NO production^67–71^. In this context, Sonkusare et al. ^6^ have reported that Ca^2+^ influx via TRPV4 activation occurs independently of NO production. Our findings from ECs exposed to GSK treatment reveal a notable decrease in NO production and concurrent reduction in hyperpolarization following prolonged exposure to L-NAME (**Supplemental Figure 7**). This observation aligns with the established role of eNOS and NO production in endothelial hyperpolarization^66^. Furthermore, our in vivo data indicate that prolonged inhibition of NO production (one week) affects the vascular response in vessels pre-treated with PE and upon GSK and ACh stimulation, highlighting the critical role of NO in regulating vasomotor tone. In addition, the use of mM concentration of eNOS blockers (L-NAME) also reduced the completed EDH-relaxation component in mesenteric arterioles ^72^. Therefore, we propose that in the study by Sonkusare et al.^6^, residual NO production persists following TRPV4 activation despite the administration of NOS blockers. This residual NO might be sufficient to initiate NO/EDH signaling and subsequent vasorelaxation.

Our findings contribute to establishing and positing that two pools of NO production exist in ECs involved in mesenteric artery relaxation via TRPV4 activation. One of these pools signals through the sGC-PKG pathway in smooth muscle, while the other signals via S-nitrosylation of critical endothelium calcium-permeable channels (e.g., Cx43 hemichannels), thereby activating endothelial K_Ca_ currents and promoting endothelial hyperpolarization.

In conclusion, our study unveils the pivotal role of endothelial TRPV4/Cx43 hemichannels in orchestrating endothelial hyperpolarization through S-nitrosylation, thereby enhancing EDH generation. This novel insight underscores the intricate interplay between TRPV4/Cx43 hemichannels and NO signaling pathways, highlighting their profound impact on endothelium electrical behavior and vascular tone regulation.

## SUPPLEMENTAL FIGURES

**Supplemental Figure 1:** Validation of Cx45 KO cell line. **A**) Sanger sequencing of Cx45 from Wild-type HeLa cells (top) and the clone # 18 used in this work. The mutation induced by CRISPR (Synthego) induced the deletion of 2 nucleotides causing a frameshift in the Cx45 sequence (asterisk). The guide sequence used for guide RNA synthesis is shown as a horizontal black line. The vertical dotted line represents the cut site. **B**) Western blot of HeLa lysates from 4 different monoclonal populations and two different batches of Wild-type HeLa cells. Clone 18 was transiently transfected with rat Cx45 as a positive control for the Cx45 antibody and to demonstrate transfection efficiency in this clone. Lysates from *Xenopus laevis* oocytes were also used as control. Oocytes were injected with RNA for Cx45 and cell lysates were obtained 2 days after RNA injection. Non-injected oocytes were used as the negative control. **C**) Summary of ICE analysis. The predicted KO scores correlates with the signal for Cx45 detected by Western blot.

**Supplemental Figure 2.** TRPV4 controls Cx43 hemichannel activity in a heterologous expression system. Activation of Cx43 hemichannels was evaluated by assessing ethidium uptake. Connexin-free HeLa cells were transfected with human EGFP-tagged alone or in combination with human TRPV4. Dye uptake was evaluated in basal conditions and in the presence of 1 µM GSK under constant superfusion (See Methods). **(A)** Representative time courses of ethidium uptake. Each group correspond to the average of several GFP-positive cells (n=22-112) from the same coverslip. **(B)** Quantification of the ethidium uptake rate. Statistical comparisons between groups were performed using one-way ANOVA and Tukey post hoc test.

**Supplemental Figure 3:** TRPV4 decreases Cx43 expression. **A**) Representative images of the EGFP signal in cells transfected with EGFP alone or EGFP + TRPV4. Images were taken with the same exposure time (500 ms) for comparison. The quantification of EGFP signal intensity with 1 s exposure is shown in the bar graph. **B)** Representative images of the EGFP signal in cells transfected with Cx43WT-EGFP alone or Cx43WT-EGFP + TRPV4. Images were taken with the same exposure time (1s). The quantification of EGFP signal intensity is shown in the bar graph. **C)** Representative images of the EGFP signal in cells transfected with Cx43C271S-EGFP alone or Cx43C271S-EGFP + TRPV4. Images were taken with the same exposure time (1s). The quantification of EGFP signal intensity is shown in the bar graph. * = p< 0.05 by unpaired t-test. n.s.= non-significant.

**Supplemental Figure 4.** Connexin43 hemichannels do not contribute to NO production in endothelial cells following TRPV4 activation. NO production was detected using DAF-2DA in primary endothelial cell cultures after stimulation with 10 nM GSK 1016790A, preceded by pretreatment with 50 µM Gap19 and 100 µM L-NAME (a NOS blocker). The bars represent the duration of stimulation.

**Supplemental Figure 5.** Inhibition of Cx43 hemichannels by Gap19 did not affect smooth muscle function in resistance arteries. **A**) Contraction curve of phenylephrine (PE) with and without 50 µM Gap19. **B)** Smooth muscle membrane potential changes (green) observed in intact isolated arteries under control conditions and in the presence of 10 µM phenylephrine. Group comparisons were conducted using Student’s t-test. The total number of mice used in the experiments is indicated in parentheses.

**Supplemental Figure 6.** Endothelial disruption impairs vasodilation response induced by GSK or ACh. **A**) Representative traces depicting vasomotor responses induced by 10 nM GSK 1016790A, 10 µM PE, 10 µM ACh, and 10 µM NO donor DEANO in control conditions, alongside conditions where endothelium damage (ED) is present. **B)** Percentage of relaxation in vessels pre-contracted with 10 µM PE upon GSK, 10 µM ACh, and 10 µM DEANO stimulation in both control and ED conditions. Group comparisons were conducted using Student’s t-test. The total number of mice used in the experiments is indicated in parentheses.

**Supplemental Figure 7:** NO production is critical for the endothelium hyperpolarization upon TRPV4 activation. **A**) Rate of NO production in ECs under different minutes of incubation time with 100 µM LNAME. **B)** Endothelium hyperpolarization peak displayed in control conditions as well as at different minutes of incubation time with 100 µM L-NAME. Group comparisons were made using nested ANOVA. The total number of mice used in the experiments is indicated in parentheses.

**Supplemental Figure 8.** Total blots. Uncut representative blots used in Figure 2C. The dotted red line indicates the molecular weight of Cx43.

## METHODS

### Mouse Breeding

Wild-type (WT) mice were obtained from Jackson Labs, bred in our animal facility, and evaluated at 4-6 months of age. All animal studies were conducted with approval from the Institutional Animal Care and Use Committee at Rutgers New Jersey Medical School and adhered strictly to National Institutes of Health guidelines.

### Primary culture of endothelial cells

We employed primary cultures of ECs isolated from arterioles in the mesenteric vascular bed. For detailed methods, please refer to the protocol on reference^73^.

### Generation of Cx45-knockout HeLa cells

Previous studies have shown that HeLa cells endogenously express Cx45^74^. Therefore, we performed a knockout of endogenous Cx45 in HeLa cells to establish a connexin-free heterologous expression system (**Supplemental Figure 1**). A GJC1 knockout cell pool was obtained by electroporation of HeLa cells with SpCas9 and a synthetic guide RNA (sequence: AAUGCGCUGGAAACAACACC, Synthego). Clonal cell lines were generated via limiting dilution and clonal expansion using EMEM media supplemented with 10% FBS and 1% Penicillin/Streptomycin). In order to obtain genotyping clones, genomic DNA was isolated with the QuickExtract DNA Extraction Solution (Lucigen). The genomic region targeted with the guide RNA was amplified by PCR (Forward primer sequence: AGGACAAGGAAGTCTGCTGC; Reverse primer sequence: GTGGGGAAGATCTGGCTCAC) and analyzed by Sanger sequencing and Inference of CRISPR Edits (ICE) analysis (Synthego). The absence of Cx45 protein in Cx45KO clones was confirmed by Western blot and the Cx45KO clone with best proliferation rate was used for subsequent cell expansion and experiments.

### Molecular Biology

Plasmids containing the cDNAs for EGFP, human TRPV4 and human Cx43 were amplified and purified using the HiSpeed® MidiKit (Qiagen). To identify Cx43 expression/distribution, the EGFP sequence was attached to the Cx43 in the C-terminus (named Cx43WT-EGFP). To this end, EGFP was synthetized and subcloned into pcDNA3.1 plasmids containing the sequence of human Cx43. The single mutation C271S was generated by Epoch Life Science from pcDNA3.1 plasmids containing the Cx43WT-EGFP sequence. C271S mutation was confirmed by DNA sequencing.

### Dye uptake assays in HeLa cells

Cx45KO HeLa cells were seeded on glass coverslips and transfected with EGFP alone, TRPV4 + EGFP, Cx43WT-EGFP, Cx43C271S-EGFP, Cx43WT-EGFP + TRPV4, or Cx43C271S-EGFP + TRPV4 using the jetPRIME® Transfection Reagent (Genesee Scientific, CA). Co-transfections were carried out using 0.5µg for each plasmid. Cells were used 1 day after transfection, where the transfection efficiency was high (∼70%) and the number of dead cells was negligible.

Hela cells were superfused at room temperature by a gravity-dependent flow system at ∼0.5 mL/min with a working solution (composition in mM: 142 NaCl, 4 KCl, 1 CaCl_2_, 5 HEPES), adjusted to pH 7.40 and 285 mOsm. Cells were visualized using an inverted Olympus IX73 microscope and a Hamamatsu digital camera (C13440-20CU). For dye uptake recordings, cells were superfused with the working solution supplemented with 10 µM ethidium bromide for 10 min (baseline) and then with the same solution containing 1 µM GSK for 15 min. Images were acquired each 20 s (50 ms exposure), using cellSens software (Olympus). To determine expression and distribution of Cx43, we acquired snapshots for Cx43-EGFP signal before and after the experiments using the same exposure for comparison. Image J software was used for data analysis.

Dye uptake rate was analyzed in non-transfected (EGFP-negative cells) to determine endogenous ethidium uptake and transfected cells (EGFP-positive cells). For the final analysis, endogenous uptake was subtracted to show the absolute ethidium uptake induced by EGFP/TRPV4/Cx43. To quantify the net ethidium uptake induced by GSK, the dye uptake rate recorded in the presence of GSK was subtracted from the basal uptake. Dye uptake rate is shown as AU/s.

### Measurements of Intracellular Ca^2+^ Concentration

ECs were exposed to 5 μM Fluo-4AM and 0.02% (w/v) Pluronic F-127 in Tyrode solution containing 5 mM MOPS, with the following composition (mM): 118 NaCl, 5.4 KCl, 2.5 CaCl2, 1.2 KH2PO4, 1.2 MgSO4, 23.8 NaHCO3, and 11.1 glucose at pH 7.4, for 30 minutes. After washing, the cells were transferred to a temperature-controlled microscope imaging chamber maintained at 37°C on the microscope stage. We analyzed 200–300 cells per cover slip, with three cover slips used for each animal. The average value from the three cover slips was calculated to determine a single value for each experiment. Variations in fluorescence intensity were expressed as F/F0, where F represents the fluorescence observed during the recording period and F0 is the baseline fluorescence value. Fluo-4AM was prepared in DMSO and diluted to the working concentration in MOPS-buffered Tyrode solution.

### Membrane Potential Recordings

Changes in membrane potential were recorded in primary cultures of ECs and in smooth muscle cells of intact, isolated mesenteric resistance arteries (120–180 μm inner diameter) using glass pulled microelectrodes filled with 3 M KCl (pipette resistance: 30–60 MΩ) connected to a DUO 773 electrometer (World Precision Instruments, Inc., FL, USA). For smooth muscle and endothelium recordings, resistance arteries were secured on a Sylgard® surface at the bottom of a chamber containing MOPS-buffered Tyrode solution (pH 7.4). To identify the impaled cell type, the microelectrode filling solution included 10 μM dextran-FITC (MW: 3000 Da) in addition to 3 M KCl. The setup was grounded with an Ag-AgCl reference electrode immersed in the buffer solution. Successful cell impalement was confirmed by a rapid negative deflection of potential, stable membrane potential under basal conditions, and a positive deflection upon withdrawal. Membrane potential changes were recorded at 100 Hz, unless specified otherwise, using the pClamp 10 data acquisition software (World Precision Instruments Inc., FL, USA). We applied Indomethacin, a cyclooxygenase inhibitor, to minimize the impact of prostaglandins on our responses.

### Detection of S-nitrosylated proteins

S-nitrosylated proteins were isolated from primary cultured of ECs. ECs were homogenized in HEN buffer (250 mM HEPES, 1 mM EDTA, 0.1 mM Neucoproine, pH 7.7) containing protease inhibitors. Samples containing 50-100 μg protein were subjected to the biotin-switch method to pull down all S-nitrosylated proteins. The separated S-nitrosylated proteins were then analyzed using 10% SDS-PAGE and transferred onto a PVDF membrane (BioRad, Hercules). A monoclonal anti-Cx43 antibody (#C8093; 1:2,000, mouse; Sigma-Aldrich) was employed to detect Cx43 protein. Signal intensity in all Western blot analyses was quantified using ImageJ (NIH). Each treatment group consisted of samples from six independent hearts.

### Proximity Ligation Assay

The subcellular distribution and potential spatial associations of S-nitrosylation (S-NO) with Cx43, TRPV4/Cx43, eNOS/Cx43, and eNOS/TRPV4 were investigated using the Proximity Ligation Assay (PLA) from Sigma-Aldrich. ECs were initially blocked and then exposed to primary antibodies sourced from various species. Detection was facilitated using oligonucleotide-conjugated secondary antibodies following the manufacturer’s protocols. Specifically, the following antibodies were utilized: monoclonal anti-Cx43 (#C8093; 1:200; Sigma-Aldrich), Anti-S-nitrosylated (SNO-Cys) rabbit polyclonal antibody from Alpha Diagnostic (Cat. NISC11-A), Anti-eNOS/NOSIII mouse monoclonal antibody from BD Transduction (Cat. 610297), and Anti-TRPV4 rabbit polyclonal antibody from Alomone Labs (Cat. ACC-034). Each experimental group comprised samples obtained from independent cultures or mesenteric bed arterioles.

For tissue preparation, the mesenteric arteries were exposed to a 4% PFA solution for 10 minutes to fix them, followed by overnight postfixation. The tissues were then dehydrated, embedded in paraffin, sectioned (5 μm), mounted on charge-coated slides, and deparaffinized using standard procedures as previously described ^7^. Antibodies used included Anti Connexin-43 mouse (Sigma Cat. C8093) and Anti-TRPV4 rabbit (Alomone Labs Cat. ACC-034).

### Hemichannel activity

ECs were incubated in MOPS-buffered physiological saline solution (pH 7.4) containing 5 μM ethidium bromide to assess hemichannel activity a well-established method to evaluate connexin hemichannel function ^14,16,24,75,76^. The opening of Cx hemichannels was evaluated by tracking the uptake of ethidium over time. Fluorescence intensity, indicative of ethidium binding to nucleic acids, was quantified using epifluorescence microscopy with excitation at 530–550 nm and emission detection at 590 nm. Variations in fluorescence intensity were expressed as F/F0, where F represents the fluorescence observed during the recording period and F0 is the baseline fluorescence value.

### Intravital microscopy

To assess in vivo relaxation, we employed an intravital microscopy protocol as previously described ^77^. For vessel diameter detection, we utilized Dextran-70KDa-FITC, as illustrated in Figure 6A. Vasoactive drugs (e.g., GSK, PE, and ACh) were topically administered in the mesenteric bed. To block Cx43 hemichannels, we administered 10 µg/kg of Gap19-TAT via retroorbital injection ^14,16,37^. To quantify diameter changes, we employed Metamorph software.

### Chemicals

All chemicals used were of analytical grade and obtained from MilliporeSigma (Billerica, MA, USA). Specifically, N^G-nitro-L-arginine methyl ester (L-NAME), MOPS, endothelial cell growth supplement from bovine pituitary, BSA, PE, ACh, GSK 1016790A, DEANO, charybdotoxin, and apamin were sourced from MilliporeSigma.

### Statistical Analysis

Values are reported as mean ± standard error. Group comparisons were assessed using paired Student’s t-test, one-way ANOVA, or two-way ANOVA with Tukey’s post-hoc test, as applicable. For electrophysiological and Ca2+ measurements, comparisons involving cells and different treatments from the same animal were treated as technical replicates. Statistical significance for these comparisons was assessed using nested one-way ANOVA as detailed by Eisner^78^. A significance level of P < 0.05 was considered statistically significant. Each figure legend specifies the respective sample size (n) and exact P value. In the case of P < 0.0001, we have added those as 0.0001. Statistical analyses were performed using GraphPad Prism version 9.5.1 and Origin version 9.

### Author Contributions

PB designed and performed the experiments, analyzed and interpreted the data, and edited the final manuscript. PSG performed and analyzed the experiments and edited the final manuscript. PS performed experiments. PAA, performed and analyzed the experiments, AVB, designed, performed and analyzed the experiments, edited final manuscript. WND, interpreted the data, and edited the final manuscript. JEC, designed experiment, interpreted the data and edited the final manuscript. ML designed and performed the experiments, analyzed and interpreted the data, drafted the manuscript, and wrote final version. All authors reviewed and approved the final manuscript.

## Acknowledgments

This work was supported by an American Heart Association AHA Career Development Award 932684 to M.A. Lillo, AHA Research Supplement to Promote Diversity in Science 23DIVSUP1054931 to Pia Burboa, NIH grant GM 067640, GM 112415 to Annie Beuve, NIH grant R01HL 146539 to Walter Durán and NIH R01GM099490 and R21HL141170 to Jorge Contreras.

## ABREVIATIONS

ACh: Acetylcholine
ATP: Adenosine Triphosphate
β-GA: Beta-glycyrrhetinic acid
BK: Calcium-activated Potassium Channels
Ca^2+^: Calcium ions
Cx: Connexin
Cx43: Connexin 43
DEANO: Diethylamine NONOate
EC: Endothelial Cells
EDH: Endothelium-Dependent Hyperpolarization
EGFP: Enhanced Green Fluorescent Protein
eNOS: Endothelial Nitric Oxide Synthase
Et: Ethidium
IKCa: Intermediate conductance calcium-activated potassium channels
KCa: Calcium-activated potassium channels
L-NAME: NG-nitro-L-arginine methyl ester
NO: Nitric Oxide
PE: Phenylephrine
PKCα: Protein Kinase C alpha
PLA: Proximity Ligation Assay
SKCa: Small conductance calcium-activated potassium channels
TRPV4: Transient Receptor Potential Vanilloid 4

