## Supplemental figures for "Endothelial TRPV4/Cx43 Signaling Complex Regulates Vasomotor Tone in Resistance Arteries"

A

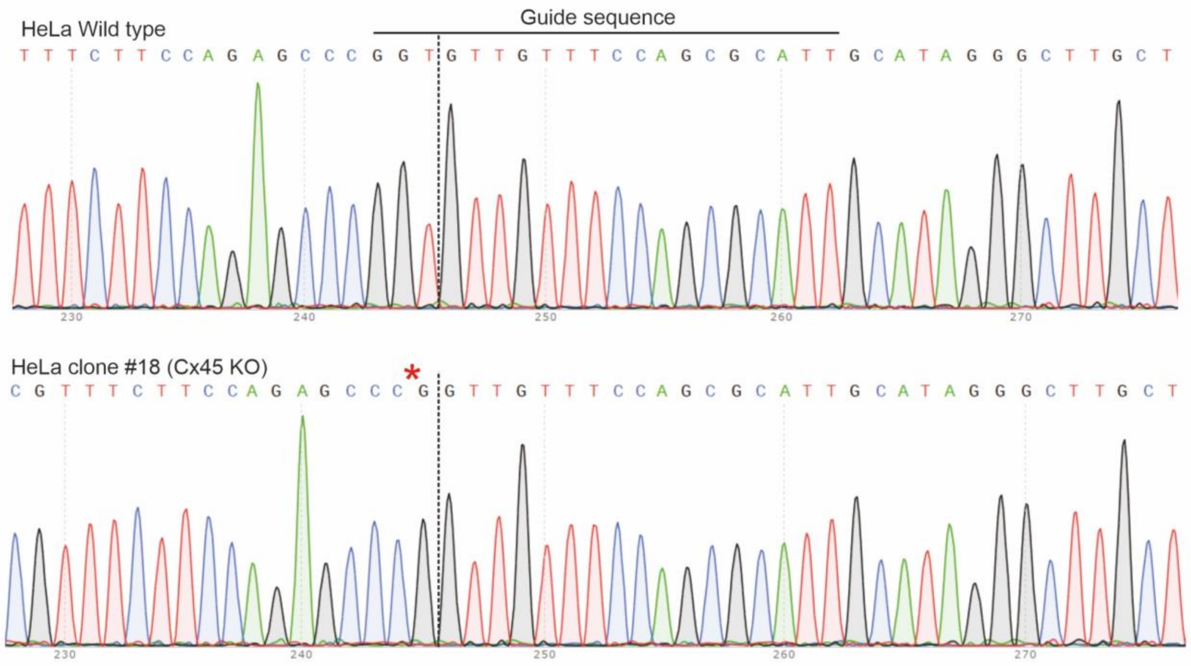


B

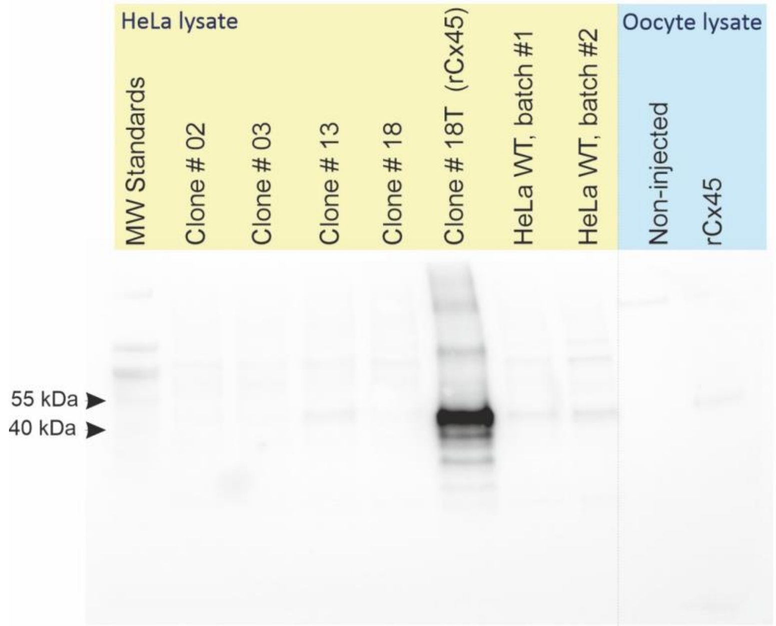


C

| | KO score by ICE analysis | Western blot signal |
|---|---|---|
| Clone # 02 | 99 | No |
| Clone # 03 | 99 | No |
| Clone # 13 | 0 | Yes |
| Clone # 18 | 100 | No |
| Wild type pool | 0 | Yes |

Supplemental Figure 1

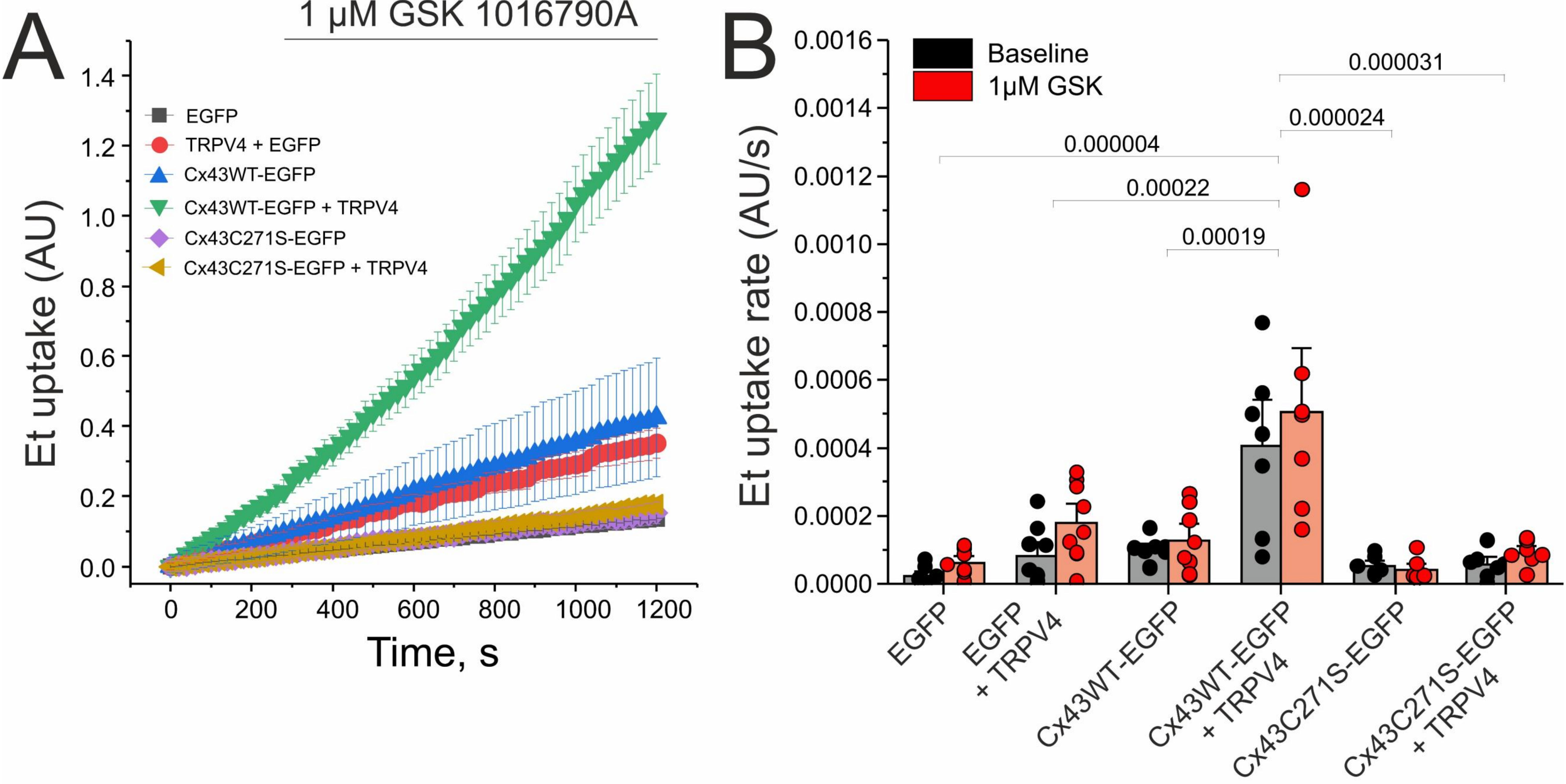


Supplemental Figure 2

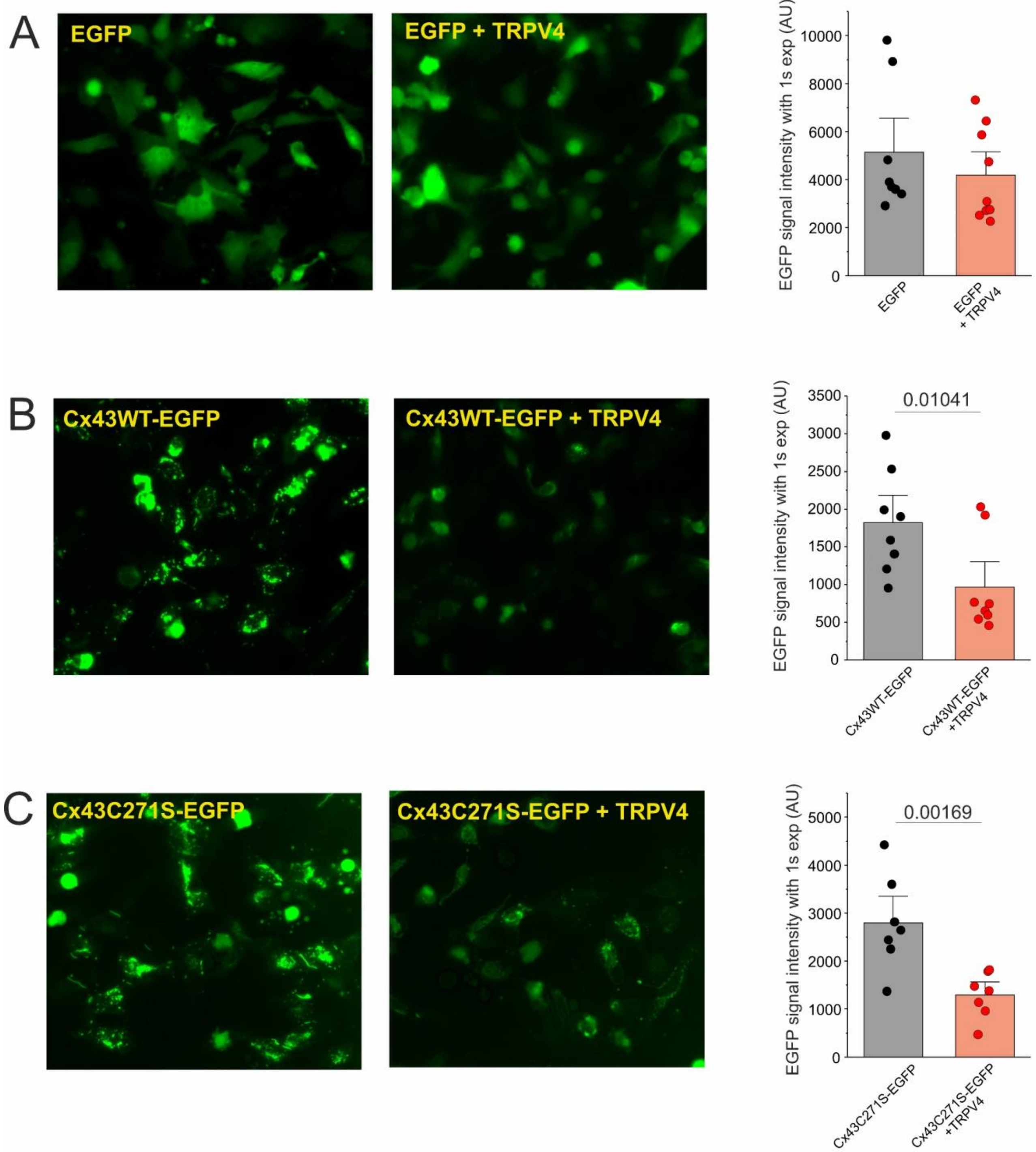


Supplemental Figure 3

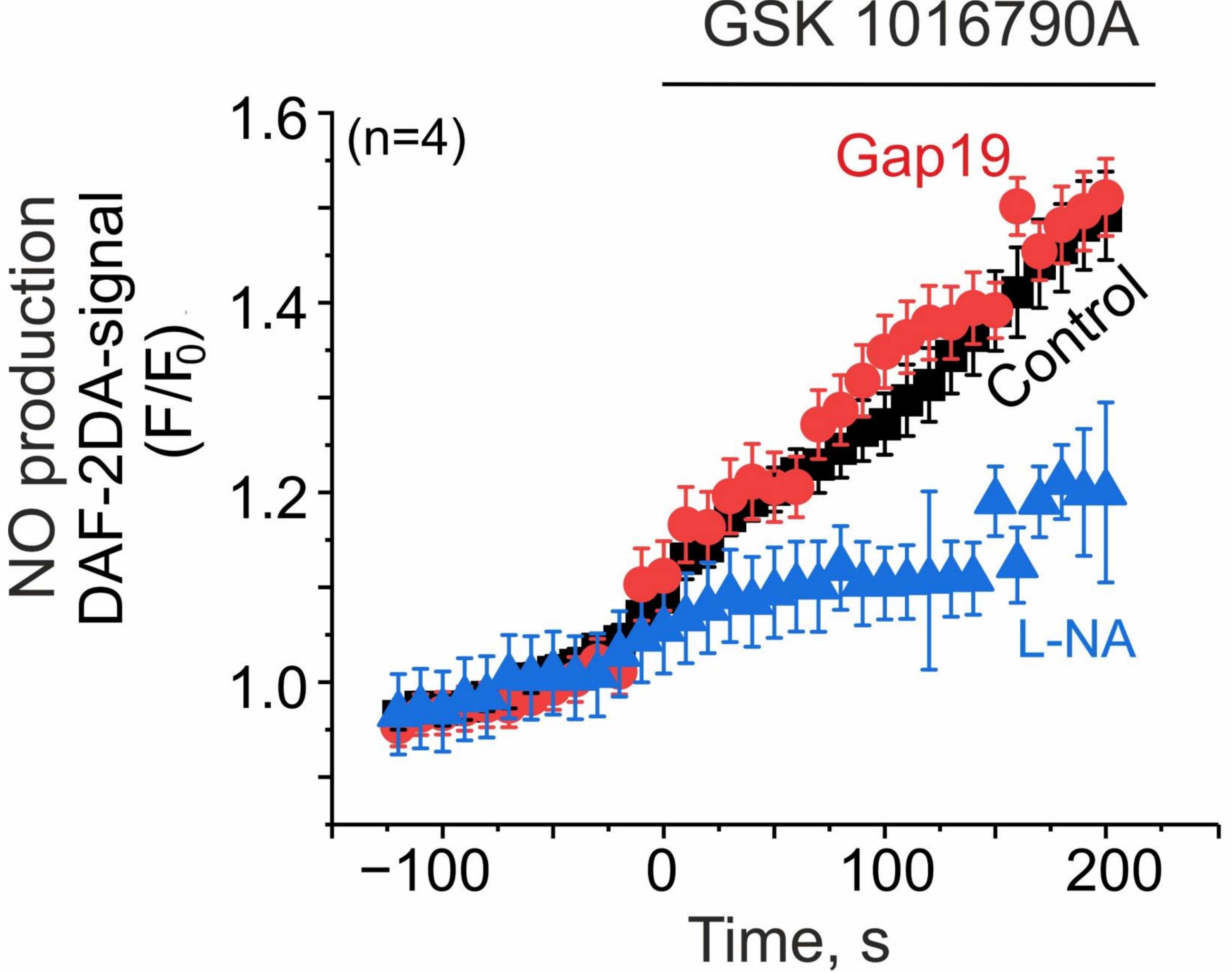


Supplemental Figure 4

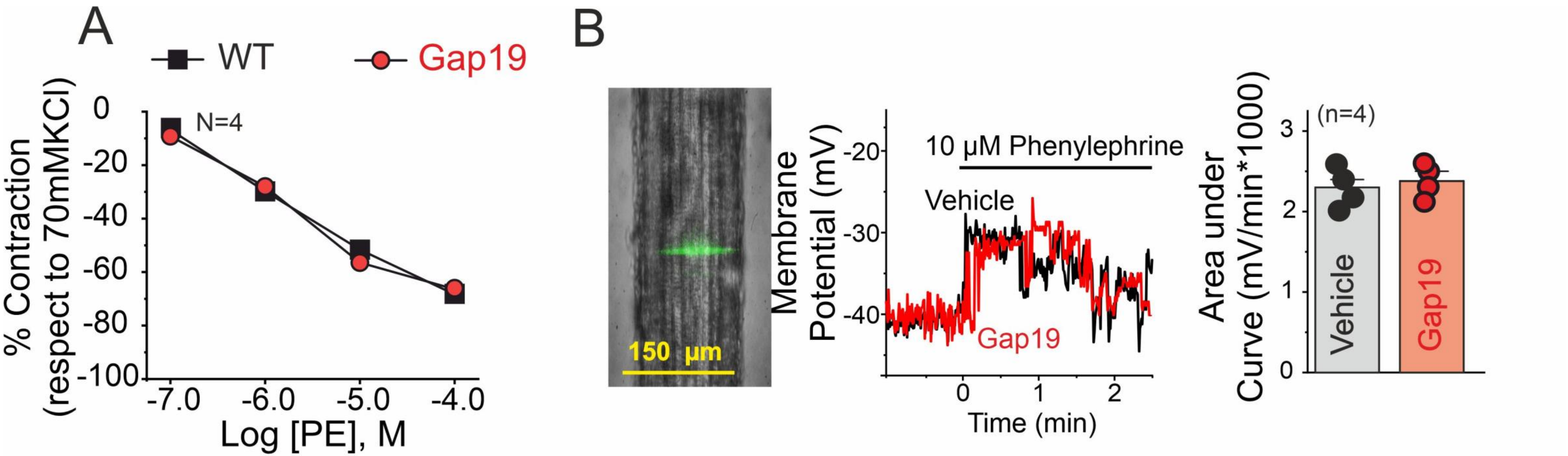


Supplemental Figure 5

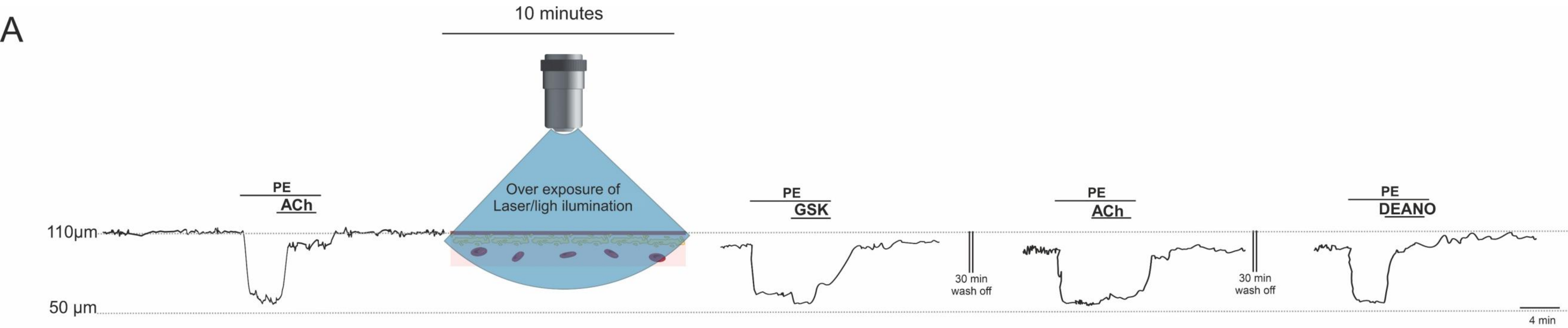


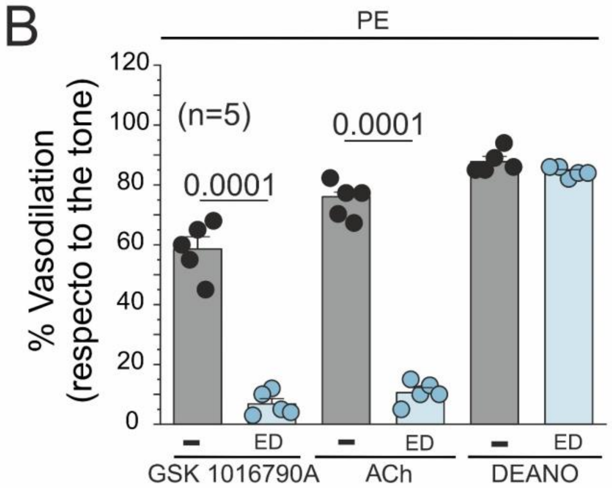


Supplemental Figure 6

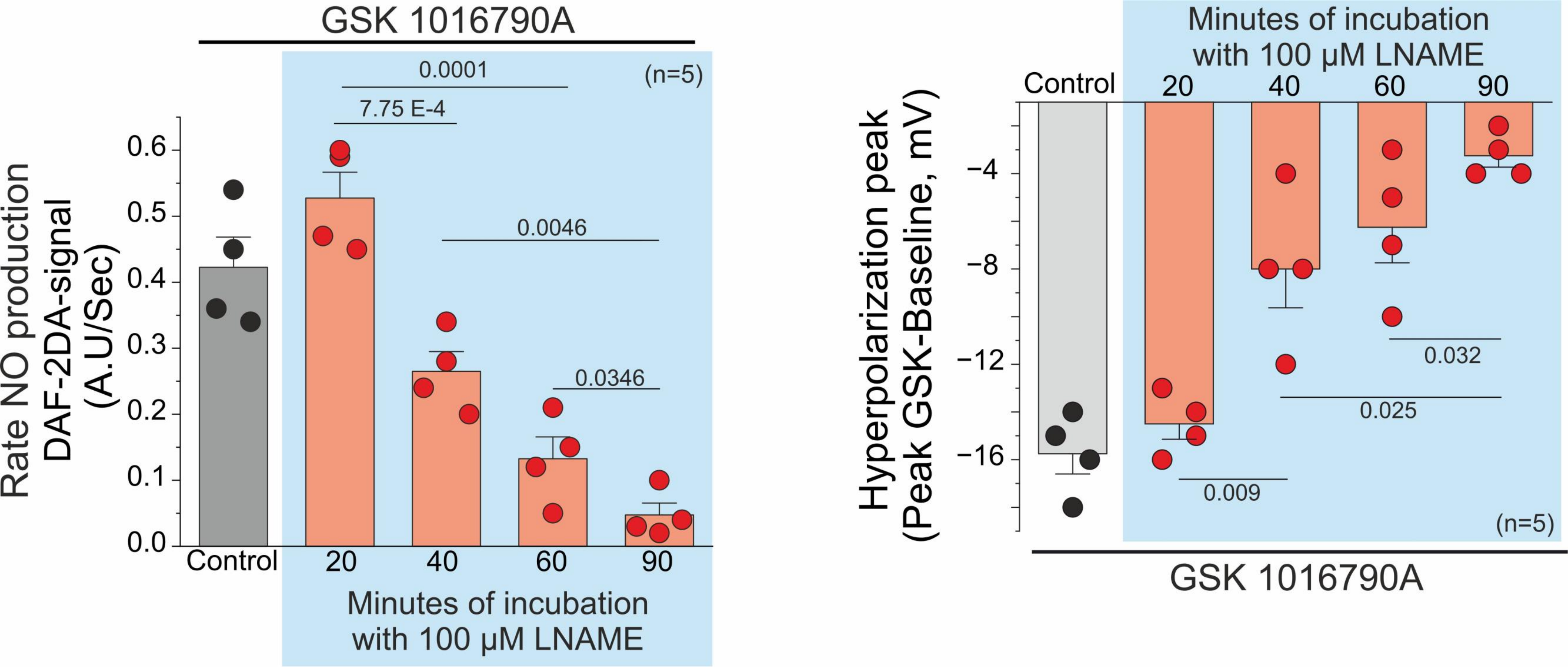


Supplemental Figure 7

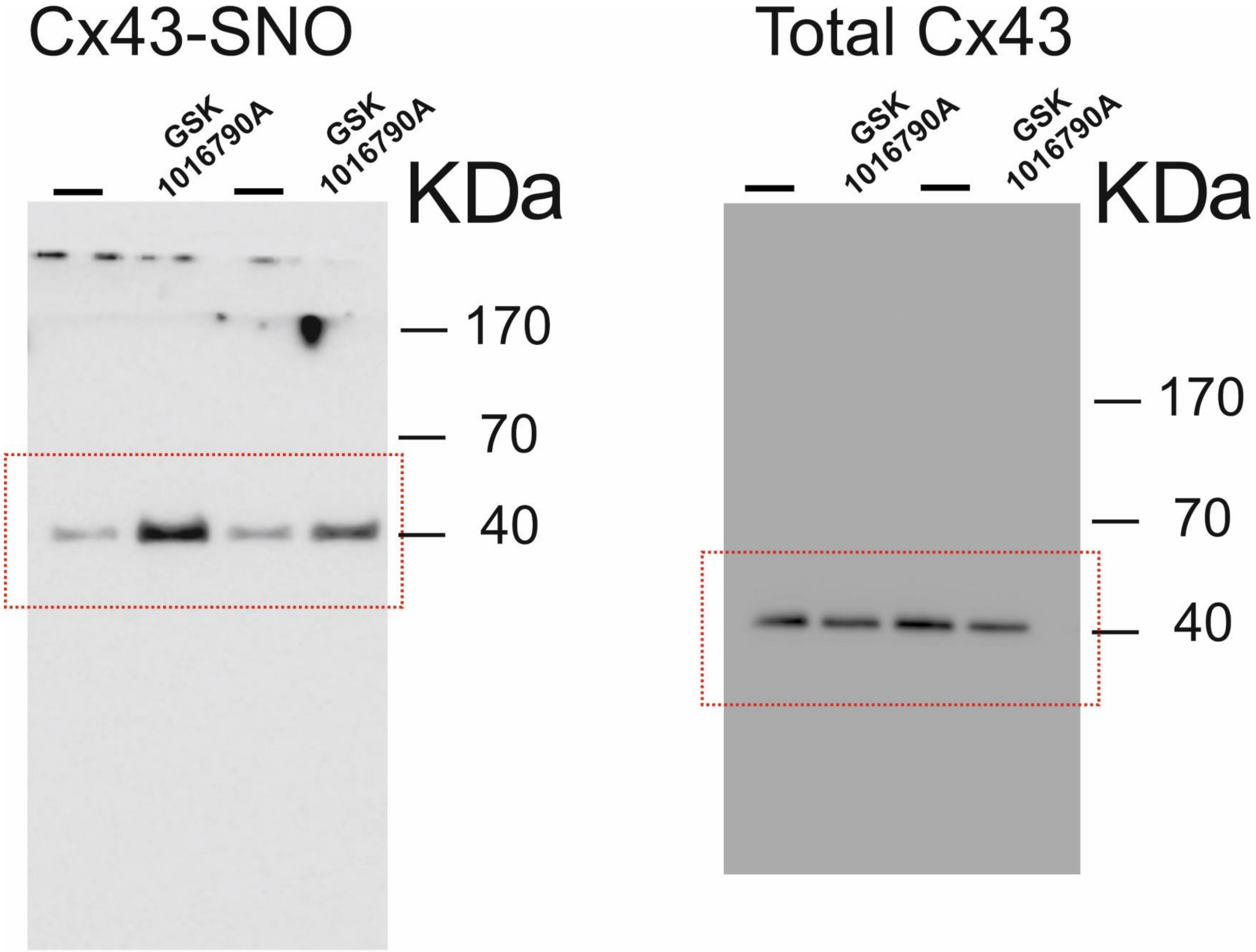

Cx43-SNO
—
GSK 1016790A
—
GSK 1016790A
KDa
170
70
40
Total Cx43
—
GSK 1016790A
—
GSK 1016790A
KDa
170
70
40
